# Differential expression of *CCD4(4B)* drives natural variation in fruit carotenoid content in strawberry (*Fragaria* spp.)

**DOI:** 10.1101/2024.07.02.601541

**Authors:** Iraida Amaya, F. Javier Roldán-Guerra, José L. Ordóñez-Díaz, Rocío Torreblanca, Henning Wagner, Waurich Veronika, Klaus Olbricht, José M. Moreno-Rojas, José F. Sánchez-Sevilla, Cristina Castillejo

## Abstract

Pigments, mainly anthocyanins and carotenoids, are important contributors to fruits’ visual appearance and nutritional properties. In strawberry (*Fragaria* spp.), the genetic and molecular mechanisms regulating fruit carotenoid biosynthesis and its natural variation remain largely unexplored. In this study, we sought to identify genomic loci contributing to variation in yellow flesh pigmentation. A major QTL, *qYellow Flesh-4B*, accounting for 82% of the total phenotypic variation was identified on *F*. × *ananassa* chromosome 4B. Following a candidate gene approach, we determined that *CCD4(4B),* a carotenoid cleavage dioxygenase (CCD), was the underlying gene. Specific polymorphisms on *CCD4(4B)* promoter region were associated with the yellow flesh phenotype and with differential regulation of *CCD4(4B)* expression during fruit ripening. Furthermore, *CCD4(4B)* expression levels were negatively correlated with violaxanthin, lutein, zeaxanthin, β-carotene and total carotenoid content. The role of CCD4(4B) in carotenoid turnover was confirmed by transient overexpression in *F.* × *ananassa* fruits, which led to a decrease in carotenoid accumulation. Notably, a −35 C>T SNP identified in *CCD4(4B)* promoter was found to be predictive for *CCD4(4B)* expression, and carotenoid content in fruits of a diverse germplasm collection, which included different octoploid *Fragaria* species. Taken together, these results provide important genetic insights into the natural variation of carotenoid content in strawberry. The High-Resolution Melting (HRM) DNA test here developed offers a fast and reliable method to predict high fruit carotenoid content, representing a useful tool for breeding projects aiming to enhance the nutritional value of this crop.

## Introduction

Modern commercial strawberry *Fragaria* × *ananassa* originated from a fortuitous cross between two wild octoploid (2*n* = 8*x* = 56) species imported to France from South and North America: *F. chiloensis* and *F. virginiana* (Duchesne, 1766). Fruits from strawberry are a rich source of bioactive compounds such as vitamin C, folate, and diverse phenolic compounds, mainly anthocyanins (Kähkönen, Hopia and Heinonen, 2001; Proteggente *et al*., 2002), which also confer the characteristic vivid red color of strawberries. As anthocyanins are quantitatively the most important pigments in strawberry, considerable effort has been devoted to understand their biosynthetic pathway and its regulation (Reviewed in (Denoyes *et al*., 2023)), with the successful identification of the causal genetic variants determining its accumulation in fruits (Castillejo *et al*., 2020). Along with flavonoid pigments, carotenoids are important contributors to fruit color, conferring hues in the yellow to red range, and also enhancing fruit nutritional value (Yuan *et al*., 2015). Certain carotenoids, such as β-carotene, serve as essential precursors to vitamin A biosynthesis, while others, like lutein and zeaxanthin, collectively known as macular pigments, are indispensable for maintaining ocular health through its ability to filter phototoxic blue light radiation and via its antioxidant activity (Arunkumar *et al*., 2018 the present, and the future). Overall, a diet rich in carotenoids is associated with a reduced risk of cancer, cardiovascular diseases, age related macular degeneration and cataract formation (Meyers *et al*., 2014; Sharoni *et al*., 2012). Strawberry is considered a fruit with a low carotenoid content (Lado, Zacarías and Rodrigo, 2016). However, natural variation has not been comprehensively studied. *F. chiloensis* yellow-tinted varieties were already described in the seventeenth century growing in the Chilean Concepción province (Ovalle, 1646). The only comparison of carotenoid composition among strawberry genotypes that has been published covered just five different *Fragaria* × *ananassa* cultivars (Zhu *et al*., 2015). Additionally, knowledge related to the carotenoid biosynthesis regulation in strawberry is also limited to three previous works performed in single *Fragaria* × *ananassa* cultivars (Zhu *et al*., 2015; García-Limones *et al*., 2008; Chen *et al*., 2023).

Chemically, carotenoids represent a large group of lipophilic isoprenoid compounds, typically featuring a C40 hydrocarbon skeleton with nine conjugated double bonds and diverse functional groups at each end of the polyene chain. Carotenoids are synthesized by all photosynthetic organisms and certain heterotrophic bacteria and fungi (Rodriguez-Concepcion *et al*., 2018). Except for very rare exceptions, animals, including humans, cannot synthesize carotenoids de novo and must rely on dietary uptake to obtain them, primarily from fruits, vegetables, and seeds. In plant cells, plastids, particularly chromoplasts in the case of flowers, fruits, and roots, are the organelles for carotenoid biosynthesis and storage (Sun *et al*., 2018). The first committed step of the carotenoid biosynthetic pathway is the condensation of two geranylgeranyl diphosphate molecules by phytoene synthase (PSY) to produce phytoene, the first C40 carotene. In non-photosynthetic tissues, PSY activity is also considered as the main limiting step of the pathway. A series of four desaturation (dehydrogenation) and two isomerization reactions increases the conjugated series of double bonds, transforming the uncolored 15-cis phytoene isomer into all-trans lycopene, a pink/red-colored carotenoid. These reactions are catalyzed by four different enzymes: PDS, phytoene desaturase (PDS), 15-cis-ζ-carotene isomerase (Z-ISO), ζ-carotene desaturase (ZDS) and carotenoid isomerase (CRTISO) (Ruiz-Sola and Rodríguez-Concepción, 2012). Carotenoid biosynthesis bifurcates after lycopene to form orange α-carotene (with one ε-ring and one β-ring cyclization) and β-carotene (with two β-ring cyclizations), which initiate the α- and β-branch of the pathway, respectively. The two lycopene cyclases, ε-cyclase (LCYE/CRTL-E) and β-cyclase (LCYB/CRTL-B), catalyze the production of end-terminal ε- and β-rings respectively (Pecker *et al*., 1996; Cunningham and Gantt, 2001). Cyclized carotenes are further oxygenated by a series of hydroxylases and epoxidases, generating yellow xanthophylls such as lutein in the α-branch or zeaxanthin and violaxanthin in the β-branch. The conversion of violaxanthin into neoxanthin finalizes the core biosynthetic pathway (Cazzonelli and Pogson, 2010; Rodriguez-Concepcion *et al*., 2018; Sun *et al*., 2018).

While carotenoid composition and content exhibit relative uniformity in green leaf tissues of plant species, they vary significantly in non-green tissues (e.g. flowers or fruits) of horticultural crops, even among specimens of the same species (Yuan *et al*., 2015). Final carotenoid concentration reflects the dynamics of multiple processes, being a net result of biosynthesis, turnover, and the plastid sink strength. The multiple conjugated double bonds in the carotenoid polyene chain are the basis for their bright color and antioxidant activity, while also make them susceptible to oxidative cleavage, leading to carotenoid degradation and the production of apocarotenoids (Sun *et al*., 2018). Carotenoid breakdown can occur through nonspecific mechanisms such as photochemical oxidation, as well as oxidation by nonspecific enzymes like lipoxygenases and peroxidases. Moreover, carotenoids can undergo specific enzymatic oxidative cleavage catalyzed by carotenoid cleavage dioxygenases (CCDs), sometimes referred to as carotenoid cleavage oxygenases (CCOs). The apocarotenoids produced from these reactions include essential phytohormones such as abscisic acid and strigolactones, as well as a diverse array of plant pigments, flavors, aromas and defense compounds (Bouvier *et al*., 2005; Sun, Tadmor and Li, 2020).

In plants, CCDs are generally encoded by small gene families (Gonzalez-Jorge *et al*., 2013), further divided into two functionally different groups: 9-cis-epoxycarotenoid dioxygenases (NCEDs) that cleave 9-cis-violaxanthin and 9-cis-neoxanthin into xanthoxin for ABA biosynthesis, and CCDs (CCD1, 4, 7 and 8) which exhibit different substrate specificities and cleavage sites to produce a large number of apocarotenoids (Sun, Tadmor and Li, 2020). Neither CCD7 and CCD8, involved in the generation of the apocarotenoid hormone strigolactone, nor NCEDs activities are expected to dramatically affect carotenoid levels in plants (Yuan *et al*., 2015). In contrast, CCD1 and CCD4 have been associated with carotenoid homeostasis and the production of apocarotenoid volatiles contributing to aroma and flavor (Sun, Tadmor and Li, 2020; Gonzalez-Jorge *et al*., 2013). Interestingly, CCD1 has cytosolic localization (Simkin *et al*., 2004; Auldridge *et al*., 2006; Rubio *et al*., 2008), what would prevent it from having direct access to the carotenoids located in the plastids. Therefore, it has been speculated that plant CCD1s could act as scavengers of the plastid-released apocarotenoids arisen through either non-enzymatic oxidative cleavage processes or enzymatic cleavage by other CCDs (Beltran and Stange, 2016). In contrast, CCD4 enzymes are located in the plastid, suggesting they may have a prominent role in carotenoid turnover. Negative correlations between CCD4 activity and total carotenoid levels have been clearly demonstrated in a number of species such as Arabidopsis, potato, *Brassica oleracea*, chrysanthemum or peach (Adami *et al*., 2013; Falchi *et al*., 2013; Campbell *et al*., 2010; Ohmiya *et al*., 2006; Han *et al*., 2019 a gene responsible for white/ yellow petal color in *B. oleracea*; Gonzalez-Jorge *et al*., 2013).

In the diploid strawberry *F. vesca*, the carotenoid dioxygenase gene family consists of seven members, named according to their respective Arabidopsis *CCD* ortholog (Wang *et al*., 2017). The NCED group includes three genes: *FveNCED1*, *FveNCED2*, and the more distantly related *FveNCED6*. Expression levels of *FveNCED2* increases rapidly during fruit development, peaking at the red stage, whereas *FveNCED1* shows a much lower expression level, with maximum expression at the pre-turning stage. *FveNCED6* could not be detected at any stage. The *CCD* gene family comprises four genes: *FveCCD1*, *FveCCD4*, *FveCCD7* and *FveCCD8*. *FveCCD1* is highly expressed throughout ripening, with a peak at the pre-turning stage followed by a continuous decline. Transcript abundance of *FveCCD4*, *FveCCD7* and *FveCCD8* is much lower and mostly decline during ripening. In *F*. × *ananassa*, induction of *FaCCD1* expression throughout fruit ripening negatively correlates with the concentration of carotenoid in fruits, especially lutein, which is cleaved by FaCCD1 in vitro (García-Limones *et al*., 2008).

Despite the limited information available regarding the diversity in carotenoid content and composition in strawberry fruit, yellow strawberry fruits have been described long ago not only for *F. chiloensis* but also for *F. vesca*, as for example the ‘Yellow wonder’ accession. The objective of this study was to identify the genetic factors controlling this variation, and the foundation was the observation of a yellow flesh phenotype segregating within the F_2_ lines of the SS×FcL mapping population previously characterized for white/red fruit color variation (Castillejo *et al*., 2020). We hypothesized that the prevailing presence of anthocyanins in strawberry fruits could be masking natural variation in yellow pigmentation and decided to investigate its chemical nature and the molecular mechanisms controlling it. Through QTL mapping, we successfully identified a major QTL, here named *qYellow Flesh-4B*, in which *CCD4(4B)* was further identified as the underlying gene. Two allelic variants of *CCD4(4B)* showing different expression levels in ripe fruits were identified in the mapping population. Notably, a negative correlation was observed between *CCD4(4B)* expression and fruit carotenoid content, particularly with the xanthophylls violaxanthin and lutein. A High-Resolution Melting (HRM) marker was developed to discriminate between the two *CCD4(4B)* alleles, which exhibited predictive capability for carotenoid content across a diverse collection of *F. virginiana*, *F. chiloensis* and *F.* × *ananassa* germplasm. Qualitative and quantitative analysis of carotenoid content in contrasting genotypes from the diverse collection, revealed significant natural variation, providing valuable insights for dietary recommendations. Additionally, the marker developed will allow the selection of favorable alleles determining higher carotenoid content in breeding programs, or facilitate its introgression from wild relatives, offering an effective strategy to augment the nutritional value of strawberry fruit.

## Results and Discussion

### Genetic mapping of the ‘Yellow Flesh’ trait reveals a single major locus in the octoploid strawberry genome

The interspecific F_2_ population SS×FcL was generated after selfing a selected F_1_ from the cross of *F.* × *ananassa* ‘Senga Sengana’ and *F. chiloensis* ssp. *lucida* USA2. We had previously used this population to map the *qFleshCol-1-2* QTL controlling internal red flesh color and the underlying gene *FaMYB10(1B)*, which controls natural variation in anthocyanin biosynthesis (Castillejo *et al*., 2020). Strikingly, all fruits lacking anthocyanins didn’t present the same coloration, showing hues that varied from white to pale-yellow, evidencing the presence of an additional colored compound that gets otherwise masked in anthocyanin rich fruits (Fig. 1A). This yellow flesh variation is also evident in other *F. chiloensis* and *F.* × *ananassa* accessions from our germplasm collection (Fig. 1B). In order to uncover the genetic control of the yellow flesh trait, we performed QTL mapping using the SS×FcL population, previously genotyped with 2,991 markers ((Castillejo *et al*., 2020); Supplementary Table S1). Ripe fruits were visually phenotyped and qualitatively scored as 1 (white) or 2 (yellow), depending on weather they showed or not yellow pigmentation. Noteworthy, the white/yellow fruit categorization throughout the manuscript merely indicates the absence/presence of yellow flesh, although internal fruit color might be red in either case due to the accumulation of anthocyanins. Fruits of all F_2_ lines accumulate anthocyanins in the epidermis with the exception of F_2_-03 that, as USA2, does not set fruit. Elevated anthocyanin accumulation in their flesh hindered phenotypic evaluation of yellow/white variation in 26 out of the 105 F_2_ lines of the population. In the remaining 78 lines, the yellow flesh segregation fitted the 3:1 expected ratio (p=0.53) for a single dominant gene, being white flesh (absence of yellow pigments) dominant over yellow flesh. QTL analysis detected a single major-effect QTL, designated *qYellow Flesh-4B,* both by Kluskal-Wallis (p<0.0001) and rMQM (LOD=28.9). The *qYellow Flesh-4B* locus was detected at the bottom of linkage group 4B (Fig. 1C), which corresponds to the top or the bottom of chromosome 4B in ‘Camarosa’ and ‘Royal Royce’ genomes, respectively. *qYellow Flesh-4B* accounted for 82% of total phenotypic variation, which may be indicative of the monogenic control of the yellow flesh trait, also in agreement with the phenotype segregation in the F_2_. Yellow coloration in strawberry fruit is likely due to the accumulation of carotenoids, as in other fruits and vegetables (Bartley and Scolnik, 1995). Genes previously studied in relation to carotenoid accumulation in strawberries, such as *FaCCD1* (García-Limones *et al*., 2008), or the lycopene cyclases *LCYB* and *LCYE* (Zhu *et al*., 2015; Chen *et al*., 2023), are located on chromosomes 7, 6, and 1 respectively. The *qYellow Flesh-4B* locus lays on a different chromosome, suggesting that, at least in the SSxFcL population, the variation in carotenoid content is not governed by these specific genes.

**Figure 1.**
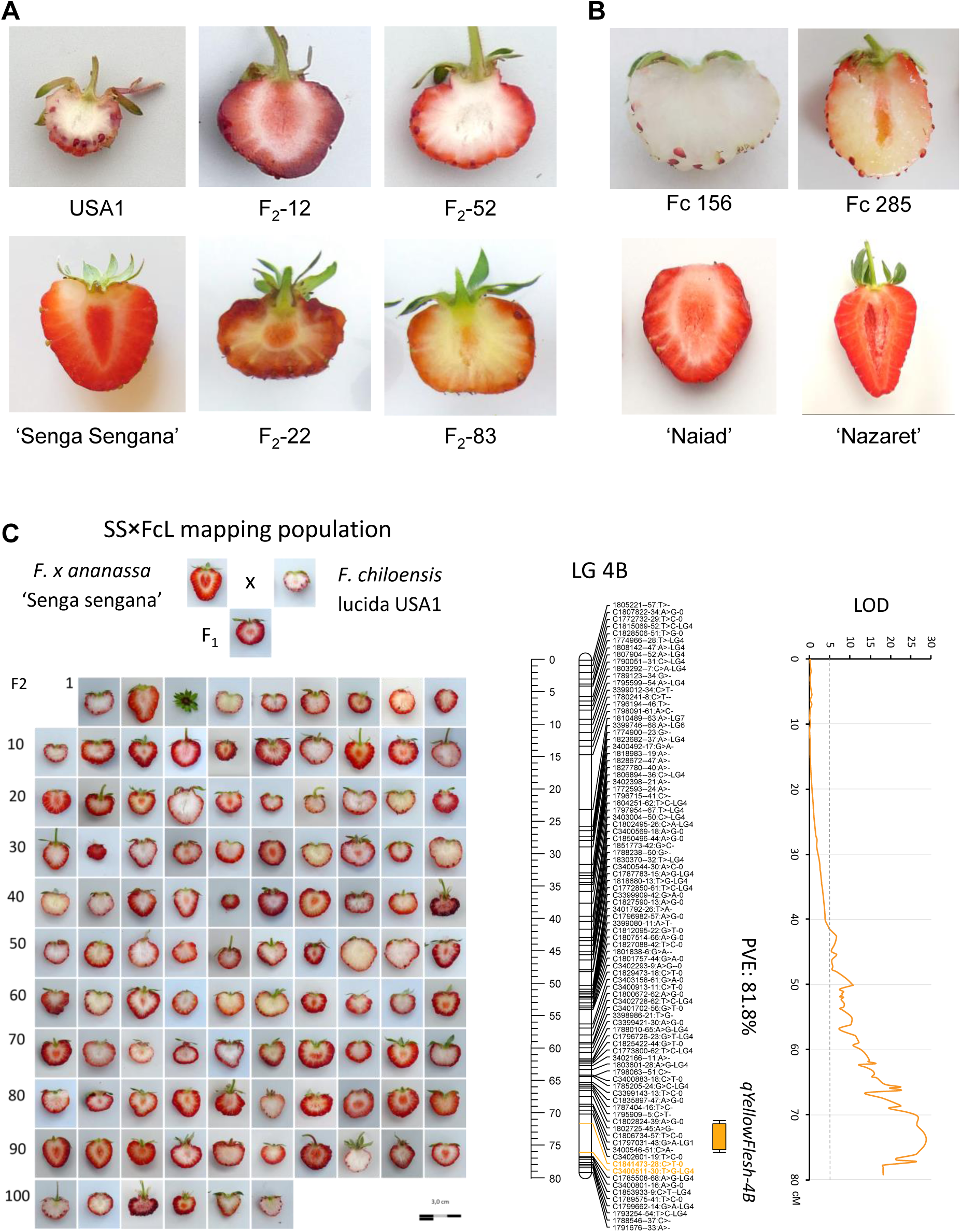
QTL Mapping of Strawberry Yellow flesh and identification of the ‘qYellow Flesh-4B’ locus. A-B) Natural variation in yellow flesh color in strawberry. A) Examples of yellow- and white-fleshed fruits among individuals of the SSxFcL population. USA1 is shown instead of USA2 (male, see Methods). USA1, F_2_-12 and F_2_-52 are white fleshed while ‘Senga sengana’, F_2_-22 and F_2_-83 are yellow fleshed. B) Two wild *F. chiloensis* and two cultivated *F.* x *ananassa* accessions differing in flesh color. C) QTL mapping using the F_2_ SSxFcL population identified the ‘qYellow Flesh-4B’ QTL on chromosome 4B as the solely contributor to fruit yellow color.

### *CCD4(4B)* is a candidate gene underlying the *qYellow Flesh-4B* locus

The *qYellow Flesh-4B* locus mapped to a reduced interval of 4.45 cM or 0.68 Mb on ‘Camarosa’ chromosome 4B, flanked by markers C1841473-28:C>T-0 and C3400511-30:T>G-LG4 (Supplementary Table S1), and encompassing 132 predicted genes (Supplementary Table S2). The corresponding chromosomal region of the diploid relative *F. vesca* contained 143 genes (Supplementary Table S3). A comprehensive exploration for candidate genes within the genomic region delimited by the QTL unveiled in both reference genomes a carotenoid cleavage dioxygenase 4 (CCD4; FxaC_14g02960 in ‘Camarosa’ and FvH4_4g34690 in *F. vesca*) as a potential contributor to the yellow flesh trait. Members of the CCD4 clade of carotenoid cleavage dioxygenases play a crucial role in carotenoid homeostasis across various plant groups (Sun *et al*., 2018). Negative correlations between CCD4 activity and total carotenoid levels have been clearly demonstrated, for example, in peach fruits and in chrysanthemum petals (Adami *et al*., 2013; Falchi *et al*., 2013; Fukamatsu *et al*., 2013; Ohmiya *et al*., 2006). Similarly, CCD4 is a negative regulator of carotenoid content in Arabidospis seeds (Gonzalez-Jorge *et al*., 2013). Consequently, the CCD4 gene in chromosome 4B, hereafter referred to as *CCD4(4B),* was selected as a candidate gene potentially involved in the regulation of white/yellow flesh color variation in strawberries. Examination of *CCD4(4B)* expression in different tissues of ‘Camarosa’ using previously obtained RNAseq data (Sánchez-Sevilla *et al*., 2017) indicated specific expression in leaf and during receptacle ripening, peaking at the white stage and drastically decreasing at the ripe stage (Supplementary Fig. S1). Orthologous CCD4s have shown species specific expression patterns, as in leaf, petal, and seed in Arabidopsis (eFP Browser at bar.utoronto.ca; (Gonzalez-Jorge *et al*., 2013), or leaf, flower, stem, and tuber in potato (Campbell *et al*., 2010). Similarly, two *CCD4* genes with contrasting expression patterns have been described in chrysanthemum, where *CCD4a* is specific of white petals while *CCD4b* is specifically expressed in leaf and stem (Ohmiya *et al*., 2006).

Sequence comparison between *CCD4(4B)* from two yellow fleshed lines (‘Senga Sengana’ and F_2_-83) and two white fleshed lines (USA2 and F_2_-52) revealed 15 polymorphic single nucleotide polymorphisms (SNPs) within the coding sequence. Among these, only one SNP resulted in a nonsynonymous substitution (p. E256D), impacting a non-conserved residue, and therefore considered unlikely to contribute to the observed phenotypic differences (Fig. 2A). In the promoter region analyzed, spanning 1 kb upstream of the *CCD4(4B)* start codon, 11 SNPs and 3 small insertions/deletions (indels) were found (Fig. 2A; Supplementary Fig. S2). Notably, *CCD4(4B)* promoter from the white fleshed lines USA2 and F_2_-52 harbored a large 432 bp insertion at −714 bp from the translation initiation codon. A query against Repbase, a repository of repetitive elements, using the CENSOR software (Kohany *et al*., 2006), identified the 432 bp insertion as a Miniature Inverted-repeat Transposable Element (MITE), exhibiting a 97% identity with DNA-5B_FV, an *F. vesca* MITE of the Mutator family. Remarkably, a similar MITE insertion in the promoter region of *CCD4b* from Citrus has been linked with enhanced gene expression, resulting in increased production of two colored apocarotenoids responsible for the red coloration of citrus fruit peel (Zheng *et al*., 2019).

**Figure 2.**
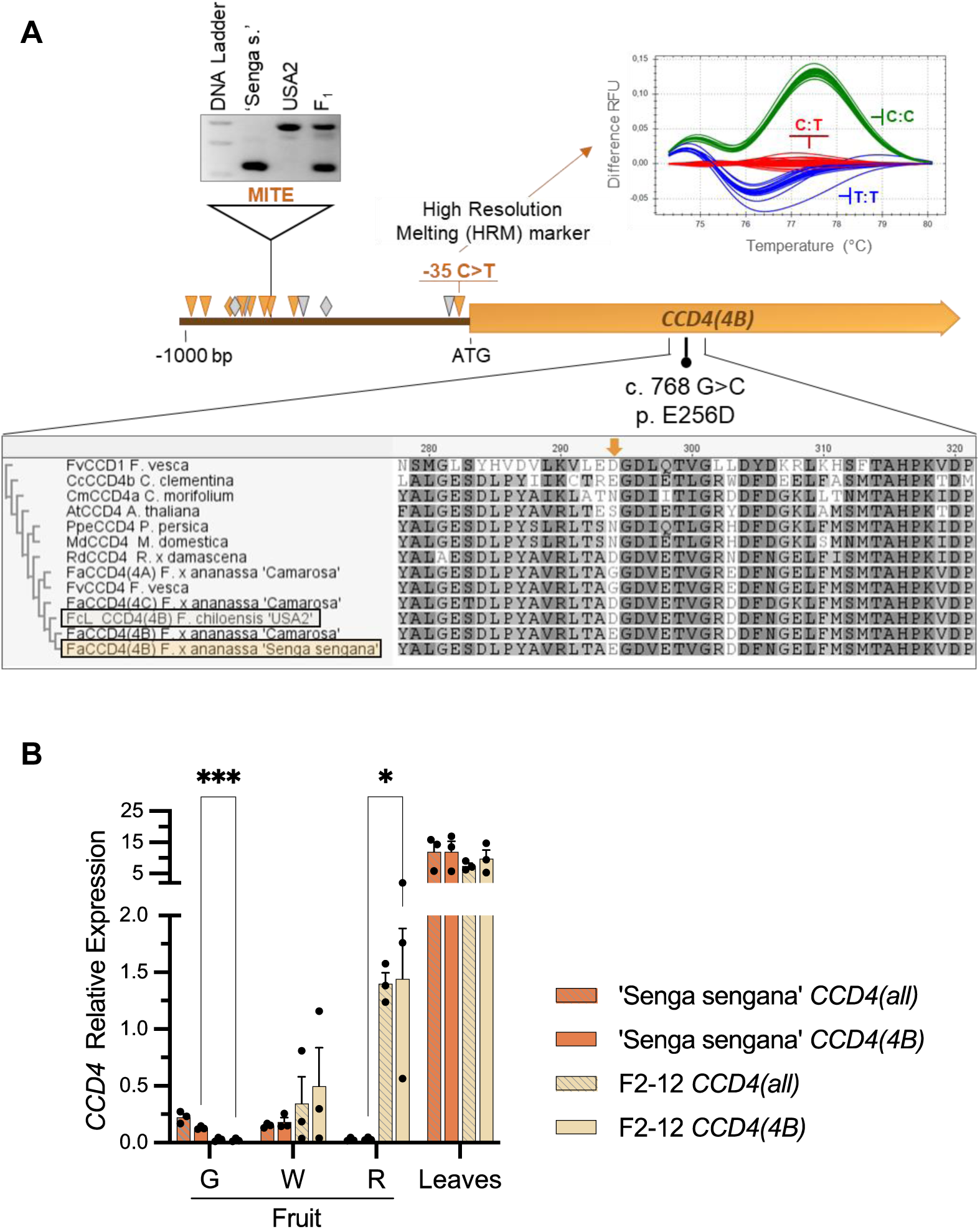
*CCD4(4B)* as Candidate Gene Underlying the ‘qYellow Flesh-4B’ QTL. A) Schematic diagram of the *CCD4(4B)* gene highlighting the polymorphisms identified between yellow- and white-fleshed individuals within the SSxFcL population. A partial alignment of CCD4 proteins is included, showing the nonsynonymous SNP (c. 768 G>C; p. E256D) found in the *CCD4(4B)* coding sequence. The two genotyping markers developed to discriminate between ‘yellow’ and ‘white’ *CCD4(4B)* alleles, one targeting the MITE insertion in the promoter region, and other an HRM marker directed to a C>T SNP located in at position −35 from the translation initiation codon has been represented. In the promoter region, triangles indicate a polymorphic SNP and diamonds, small indels. The orange ones were associated with flesh color in all accessions tested in this work, grey ones only in the SSxFcL population. B) *CCD4* expression at various stages of fruit development: green fruit (G), white fruit (W) and red fruit (R) and in leaves. ‘Senga sengana’ is the yellow-fleshed parental of the population, while F2-12 is a white-fleshed F2 sibling. Primers were designed to amplify all three *CCD4* homeolog genes (*CCD4*(all)), as well as specifically targeting the homeolog copy *CCD4(4B)*. The averages ± SEM of three biological replicates are represented. An unpaired t-test was used to compare *CCD4(4B)* expression in different tissues between the two genotypes. *P < 0.05; **P < 0.01; ***P < 0.001.

To assess the impact of the identified promoter polymorphisms on *CCD4(4B)* expression levels, RT-qPCR analysis was conducted at three stages of fruit development and ripening (green, white, and red) and in leaves. The genotypes used for the analysis were the yellow fleshed parental ‘Senga Sengana’ and the white fleshed line F_2_-12. Three homoeologous copies of the *CCD4* gene are annotated in the ‘Royal Royce’ v1.0 and ‘Camarosa’ v1.0.a1 octoploid *F.* × *ananassa* reference genomes (Hardigan *et al*., 2021a; Edger *et al*., 2019), missing the copy from chromosome 4D/4-1. In the qPCR assay, two primer pairs were employed: one targeting the three homoeologs of *CCD4* (*CCD4(4A)*, *CCD4(4B)*, and *CCD4(4C))*, and a second subgenome-specific pair directed to *CCD4(4B)*. Identical *CCD4* expression values were obtained with both primer pairs, indicating that *CCD4(4B),* located in the ‘qYellow Flesh-4B’ interval, is the dominant *CCD4* homeolog in octoploid strawberry, and the expression of the other two homeologs is negligible (Fig. 2B). Dominance of *CCD4(4B)* could also be observed in the RNAseq data from different ‘Camarosa’ tissues (Sánchez-Sevilla *et al*., 2017) although in this genotype, *CCD4(4A)* transcripts could be detected, particularly in leaves (Supplementary Fig. S1). Transcriptional profiling of *CCD4(4B)* revealed noteworthy differences between ‘Senga Sengana’ and line F_2_-12 in fruit tissue, while comparable expression levels were observed in leaves (Fig. 2B). In the yellow fleshed ‘Senga Sengana’, *CCD4(4B)* expression level remained low and predominantly decreased during ripening. However, the white fleshed line F_2_-12 showed a progressive upregulation of *CCD4(4B)* during ripening, At the red ripe stage, where the expression picked, *CCD4(4B)* levels were approximately 45 times higher than in ‘Senga Sengana’ (Fig. 2B). The *CCD4(4B)* expression rise in ripe fruits is expected to result in a higher rate of carotenoid degradation, resulting in reduced pigmentation, and therefore supporting *CCD4*(*4B)* as a promising candidate underlying the *qYellow Flesh-4B* locus. Noteworthy, previous studies in the diploid strawberry *F. vesca* ‘Yellow Wonder’, which also accumulates yellow pigment, showed that *FveCCD4* shares a similar expression pattern through fruit development with *CCD4(4B)* in ‘Camarosa’ (Supplementary Fig. S1) and ‘Senga Sengana’, the yellow fleshed parent of the SS×FcL population (Wang *et al*., 2017). This result suggests that a similar mechanism might also regulate fruit carotenoid levels in the more distantly related *F. vesca* species.

### A *CCD4(4B)* allele from *F. chiloensis* ssp. *lucida* is associated with upregulation of *CCD4(4B)* and reduced fruit carotenoid content in the SS×FcL population

To study the association between the two *CCD4(4B)* haplotypes identified segregating in the SS×FcL population and the white/yellow flesh, two PCR assays were developed to genotype the complete mapping population (Fig. 2A). One of the assays was intended to detect the MITE insertion at *CCD4(4B)_pro_* using a primer pair that flanks the insertion site. The second PCR assay was a High-Resolution Melting (HRM) marker that targets a C>T SNP at position −35 relative to the *CCD4(4B)* translation initiation codon (Fvb4-4: 1507780 *Fragaria* x *ananassa* v1.0.a1 ‘Camarosa’; chr_4B: 29120746 *Fragaria* x *ananassa* ‘Royal Royce’ v1.0). In the four sequenced *CCD4(4B)_pro_* fragments, white accessions harbored a T nucleotide at the −35 SNP (*CCD4(4B)*^T^) and the −714 MITE insertion, whereas the yellow fleshed were *CCD4(4B)*^C^ and did not carry the MITE insertion (Supplementary Fig. S2, Supplementary Table S4). As expected, genotypes of the whole SS×FcL population also showed a complete association between the two markers. Additionally, in 79 (97.5%) out of the 81 lines (including parental, F_1_ and F_2_ lines) in which the phenotype could be recorded, white flesh was associated with having at least one copy of the dominant MITE/*CCD4(4B)*^T^ haplotype, whereas yellow fleshed lines were homozygous for the allele without the MITE insertion and *CCD4(4B)*^C^ (Supplementary Fig. S3, Supplementary Table S4). In the two lines where the genotype did not predict fruit color, a phenotyping error cannot be discarded. Alternatively, it might also represent the variance not explained by the *qYellow flesh-4B* locus.

To further investigate the role of CCD4(4B) in controlling yellow flesh in strawberry, linkage between the identified polymorphisms and *CCD4(4B)* expression levels was analyzed by RT-qPCR. As shown in Fig. 2B, the largest change in *CCD4(4B)* expression along ripening between yellow fleshed ‘Senga Sengana’ and white fleshed F_2_-12 was observed at the red ripe stage. At this stage, a sharp upregulation of *CCD4(4B)* was observed exclusively in F_2_-12. Therefore, this ripening stage was selected to analyze the expression of the two *CCD4(4B)* alleles in eight additional lines with contrasting phenotype. The parental USA2, genetically white fleshed, could not be included in the analysis as it is a male plant and do not produce fruits. Instead, the trioecious relative USA1, which is a female and fertile plant, was analyzed. Consistent with the results observed in F_2_-12, expression of *CCD4(4B)* was strongly induced in ripe fruit of lines carrying the MITE*/CCD4(4B)*^T^ haplotype compared to those with the no-MITE/*CCD4(4B)*^C^ haplotype (Fig. 3A).

**Figure 3.**
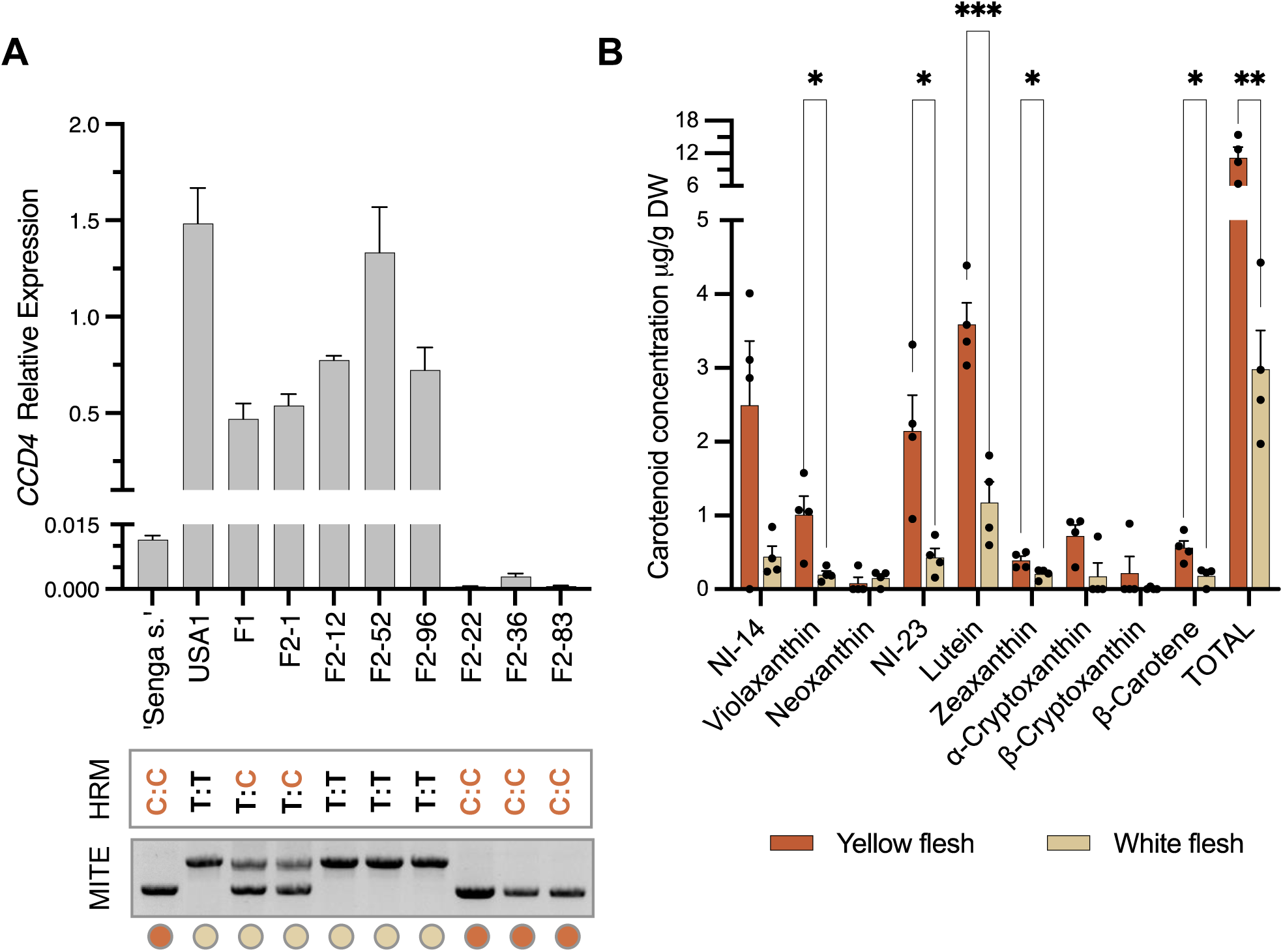
A *CCD4(4B)* Allele Correlates with Increased Fruit Carotenoid Content in the SS×FcL Population. **A)** Expression of *CCD4(4B)* in ripe fruit from different lines of the SS×FcL population. Average ± SEM of three biological replicates are represented. The genotype and phenotype (orange circle = yellow flesh; clear circle = white flesh) of each line are indicated below the graphs. The dominant MITE/*CCD4(4B)*^T^ haplotype is associated with higher *CCD4(4B)* expression levels. **B)** Carotenoid profiles were assessed by HPLC in ripe fruit from four lines of each phenotype, with three biological replicates per line. Yellow fleshed lines: ‘Senga sengana’, F_2_-22, F_2_-36, F_2_-83. White fleshed lines: USA1, F_2_-1, F_2_-12 and F_2_-52. Average values from the four lines with the same phenotype ± SEM were plotted. Two carotenoid compounds, NI-14 and NI-23, could not be identified. Multiple unpaired t-test were conducted to compare carotenoid concentrations between the two phenotypes. *P < 0.05; **P < 0.01; ***P < 0.001.

CCD4 catalyze the oxidative cleavage of carotenoids into apocarotenoids. Therefore, fruits with increased CCD4 activity are expected to have lower carotenoid content. To test the effect of differential *CCD4(4B)* expression on carotenoid accumulation in fruit, carotenoid content was determined by HPLC in ripe fruit from four yellow fleshed lines (and consequently low *CCD4(4B)* expression: ‘Senga Sengana’, F_2_-22, 36 and 83) and four white fleshed lines (and high *CCD4(4B)* expression: USA1, F_2_-1, 12 and 52). As predicted, the average total carotenoid content in the four yellow fleshed lines was significantly higher than in white fleshed lines, 11.23 ±1.91 μg/g DW vs. 2.99 ±0.52 μg/g DW (Fig. 3B, Supplementary Table S5). Regarding carotenoid profiles, nine different carotenoids were detected and compared between yellow and white fleshed samples. The most abundant one was lutein, which constituted on average 32-39% of the total amount of carotenoids, followed by, in order of concentration, violaxanthin, α-cryptoxanthin, β-carotene, zeaxanthin, β-cryptoxanthin and neoxanthin. Two additional highly abundant but unidentified carotenoid compounds were also detected, non-identified (NI)-14 and NI-23, with retention times of 14.4 min and 22.13/23.59 min, respectively. Levels of phytoene, lycopene, and α-carotene were below the detection limit in these samples, suggesting that these carotenoids might not accumulate in strawberry fruit. Our results agree with previous studies also reporting lutein as the predominant carotenoid compound in strawberry fruit (García-Limones *et al*., 2008; Zhu *et al*., 2015). In essence, the higher concentration of β-carotene and xanthophylls in yellow fleshed fruits is consistent with their flesh color phenotype and supports the role of *CCD4(4B)* in the control of carotenoid accumulation in strawberry fruit, specifically by modulation of its expression levels.

### Functional validation of *FaCCD4(4B)* role in carotenoid homeostasis in strawberry

The validation of *FaCCD4(4B)* function in the regulation of fruit carotenoid content, was conducted by transient *CCD4(4B)* overexpression in *F.* × *ananassa* ‘Rociera’ fruits. We decided to use the Spanish cultivar ‘Rociera’ instead of other cultivated accessions because its yellow flesh is more apparent than in other accessions as ‘Camarosa’ and more productive in Spain than the German cultivar ‘Senga Sengana’. The *CCD4(4B)* CDS was amplified from the white accession F_2_-52 and placed under the control of the constitutive CaMV35S promoter for agroinfiltration. A parallel set of fruits served as control and was agroinfiltrated with the pBI-GUS plasmid (GUS control). ‘Rociera’ cultivar, as described in the next section, is homozygous for *CCD4(4B)*^C^ and accordingly fruits barely express *CCD4(4B)* at the ripe stage (Fig. 4A). In fruits agroinfiltrated with *35S:CCD4(4B)*, there was a substantial increase in *CCD4(4B)* expression, reaching 120-225 times higher levels compared to the GUS control (Fig. 4A). This overexpression of *CCD4(4B)* resulted in a significant 45% reduction of total fruit carotenoid content (Fig. 4B, Supplementary Table S6), confirming the involvement of *CCD4(4B)* in carotenoid turnover in strawberry fruits.

**Figure 4.**
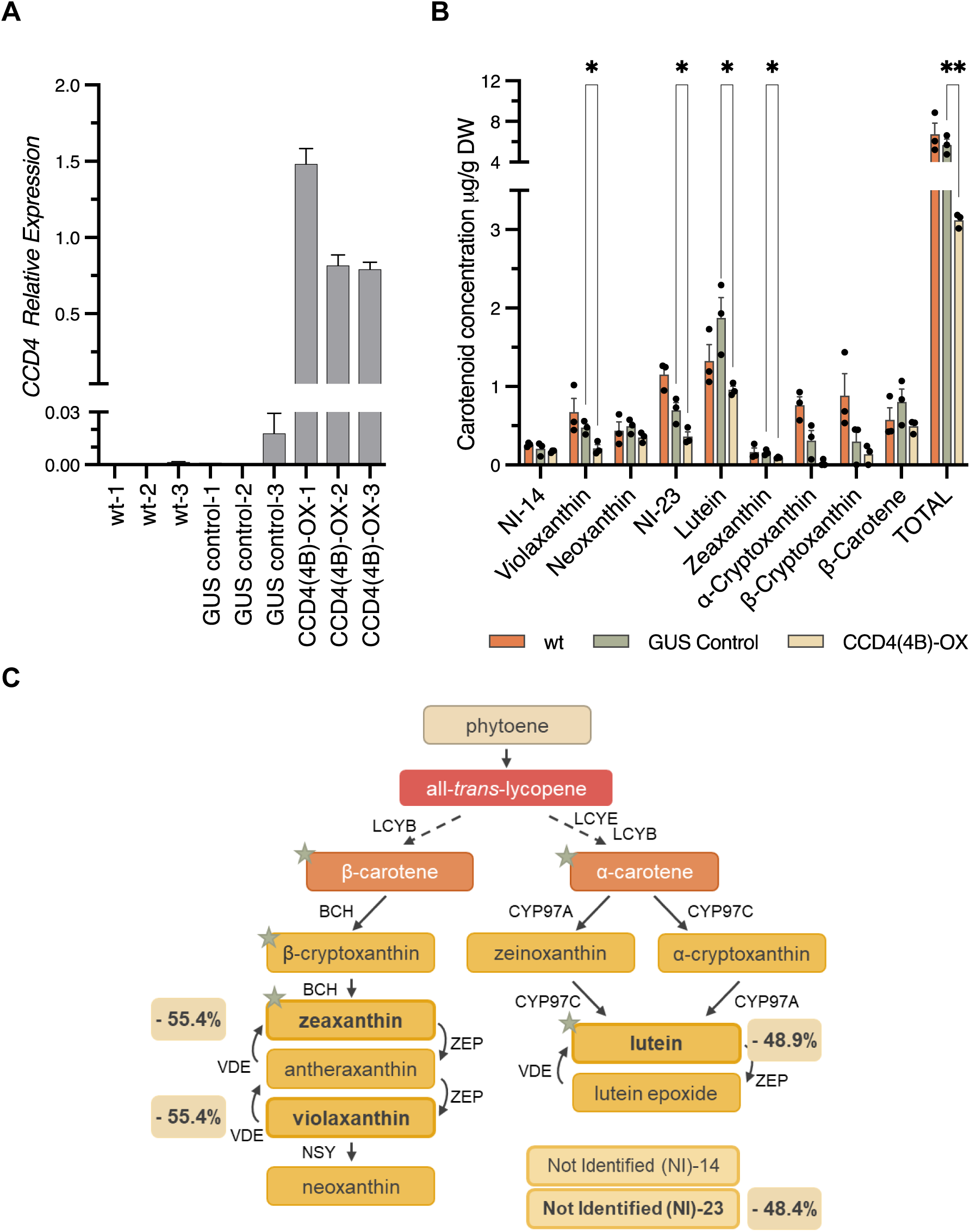
Functional validation of *CCD4(4B)* Role in the Regulation of Fruit Carotenoid Content. **A)** Expression of *CCD4(4B)* in ripe untransformed (WT) strawberry fruits (*F*. × *ananassa* ‘Rociera’), and following Agrobacterium-mediated transient transformation. GUS control: control fruits transformed with vector pBI-GUS and the P19 suppressor of gene silencing. CCD4(4B)-OX: fruits co-transformed with *35S*:*CCD4(4B)* and P19. **B)** HPLC carotenoid profiles in agroinfiltrated fruits. The average ± SEM of three independent biological replicates are represented. Multiple unpaired t-test were conducted to compare carotenoid concentration values between GUS control and CCD4(4B)-OX agroinfiltrated fruits. *P < 0.05; **P < 0.01. **C)** Schematic diagram of the carotenoid biosynthesis pathway. Star symbols indicate putative CCD4 substrates based on existing literature. Xanthophyll carotenoids with statistically significant differences in *35S:CCD4(4B)* fruits, along with the corresponding percentage decrease, are highlighted in bold.

CCD4 enzymes exhibit distinct substrate specificity and regioselectivity across plant species, contributing to the diverse array of apocarotenoid volatiles and pigments (reviewed in (Zheng, Yang and Al-Babili, 2021)). Known substrates for CCD4s in various species include α-carotene, β-carotene, β-cryptoxanthin, lutein and zeaxanthin (Rodriguez-Concepcion *et al*., 2018). In our analysis, the carotenoids NI-23, lutein, zeaxanthin and violaxanthin showed a significant decline in *35S:CCD4(4B)* overexpressing lines compared to the GUS control (Fig. 4C). Notably, the same compounds were found to be significantly different in contrasting individuals from the SS×FcL population, with the only difference that in the SS×FcL population, β-carotene also exhibited significant differences (Fig. 3B). These findings suggest that at least the two isomers zeaxanthin and lutein, also known as macular pigments, may serve as natural substrates of strawberry CCD4(4B). While violaxanthin concentration also shows a significant 55% reduction, this decrease may be induced by the lower content of the upstream xanthophyll zeaxanthin. Alternatively, violaxanthin might be also recognized as a substrate for strawberry CCD4(4B), as shown for potato (*Solanum tuberosum*) CCD4 (Campbell *et al*., 2010).

### The −35 C/T SNP at *CCD4(4B)_pro_* predicts *CCD4(4B)* expression level and carotenoid content in a diverse multi-species strawberry collection

We have shown that in the SS×FcL population, *CCD4(4B)* is a key genetic factor controlling the observed natural variation in yellow flesh pigmentation. DNA polymorphisms in *CCD4(4B)_pro_* were associated with differential *CCD4(4B)* expression, which has a direct impact on fruit carotenoid accumulation. To assess if this genotype-phenotype association also operates beyond the SS×FcL population, we expanded our analysis to a diverse collection of 222 octoploid *Fragaria* accessions from the IFAPA germplasm collection (ESP138). This selection encompasses six *F. virginiana*, twenty-four *F. chiloensis*, and 178 *F.* × *ananassa* cultivars and hybrids, representing over 125 years of breeding efforts and a wide geographic diversity.

Initially, *CCD4(4B)_pro_* was PCR amplified and sequenced from a subset of eleven *F. chiloensis* and *F.* x *ananassa* genotypes (Supplementary Fig. S2). Remarkably, the MITE insertion previously detected in SS×FcL white fleshed individuals was found in only one newly genotyped accession, *F. chiloensis* (Fc) 595 (Supplementary Fig. S2, Supplementary Table S7) the only representative of the *lucida* subspecies in the collection alongside USA1 and USA2. Only eight SNPs and one small indel, from the eleven and three, respectively identified between the yellow and white *CCD4(4B)* haplotypes, were associated with flesh color phenotype in this extended collection (Fig. 2A). Among them, the −35 C>T SNP at *CCD4(4B)_pro_*, which exhibited higher polymorphism among the evaluated accessions than the MITE insertion (Supplementary Table S7). Discrepancies between the MITE and HRM markers indicate they are not linked beyond the SS×FcL population, implying that at least one of them may not be predictive for fruit carotenoid content.

Genotyping with the −35 C>T SNP HRM marker revealed distinct allelic frequencies among the cultivated *F.* × *ananassa* accessions and the two octoploid species’ ancestors, *F. chiloensis* and *F. virginiana*. All *F. virginiana* genotypes were *CCD4(4B)*^C^ homozygous, whereas *F. chiloensis* accessions predominantly featured the *CCD4(4B)*^T^ allele, with seventeen out of the twenty-four genotypes being *CCD4(4B)*^T^ homozygous, four heterozygous *CCD4(4B)*^T^:*CCD4(4B)*^C^ and the remaining three *CCD4(4B)*^C^ homozygous (Supplementary Table S8). Strikingly, the 163 *F.* × *ananassa* cultivars were almost exclusively homozygous for *CCD4(4B)*^C^, with only 11 accessions being heterozygous *CCD4(4B)*^T^:*CCD4(4B)*^C^ and no *CCD4(4B)*^T^ homozygote found (Supplementary Table S9). As expected, *CCD4(4B)*^T^ allele frequency was higher in *F. chiloensis* introgressed hybrids (Supplementary Table S9).

To gain further insight into the association between each allelic variant and fruit carotenoid content, *CCD4(4B)* expression was analyzed in selected *CCD4(4B)*^C^ and *CCD4(4B)*^T^ accessions from the three species. Among *F. virginiana accessions,* only Fv742 produced fruits under our growing conditions and was included in the analysis. As observed in the SS×FcL population, *CCD4(4B)* expression was markedly induced at the ripe stage in those genotypes containing at least one copy of the *CCD4(4B)*^T^ allele. Interestingly, expression levels were not associated with the presence of the MITE insertion (Fig. 5A and 5B), as only Fc595 had the MITE insertion, indicating that the transposon is not responsible of *CCD4(4B)* upregulation in these diverse accessions. Alignment of the 1kb *CCD4(4B)_pro_* region sequenced from the eleven *F. chiloensis* and *F.* × *ananassa* genotypes revealed that, in addition to the −35 C>T SNP, there were seven other polymorphic SNPs and one small indel associated with differential *CCD4(4B)* expression (Supplementary Table S7, Supplementary Fig. S2). However, further functional studies would be required to verify which, if any, of the identified polymorphisms could be the causal mutation in *CCD4(4B)* behind natural variation in yellow flesh. It can also not be ruled out that different polymorphisms could be the causal mutation in different *Fragaria* accessions, as previously described for other traits such as anthocyanins and *MYB10* (Castillejo *et al*., 2020).

**Figure 5.**
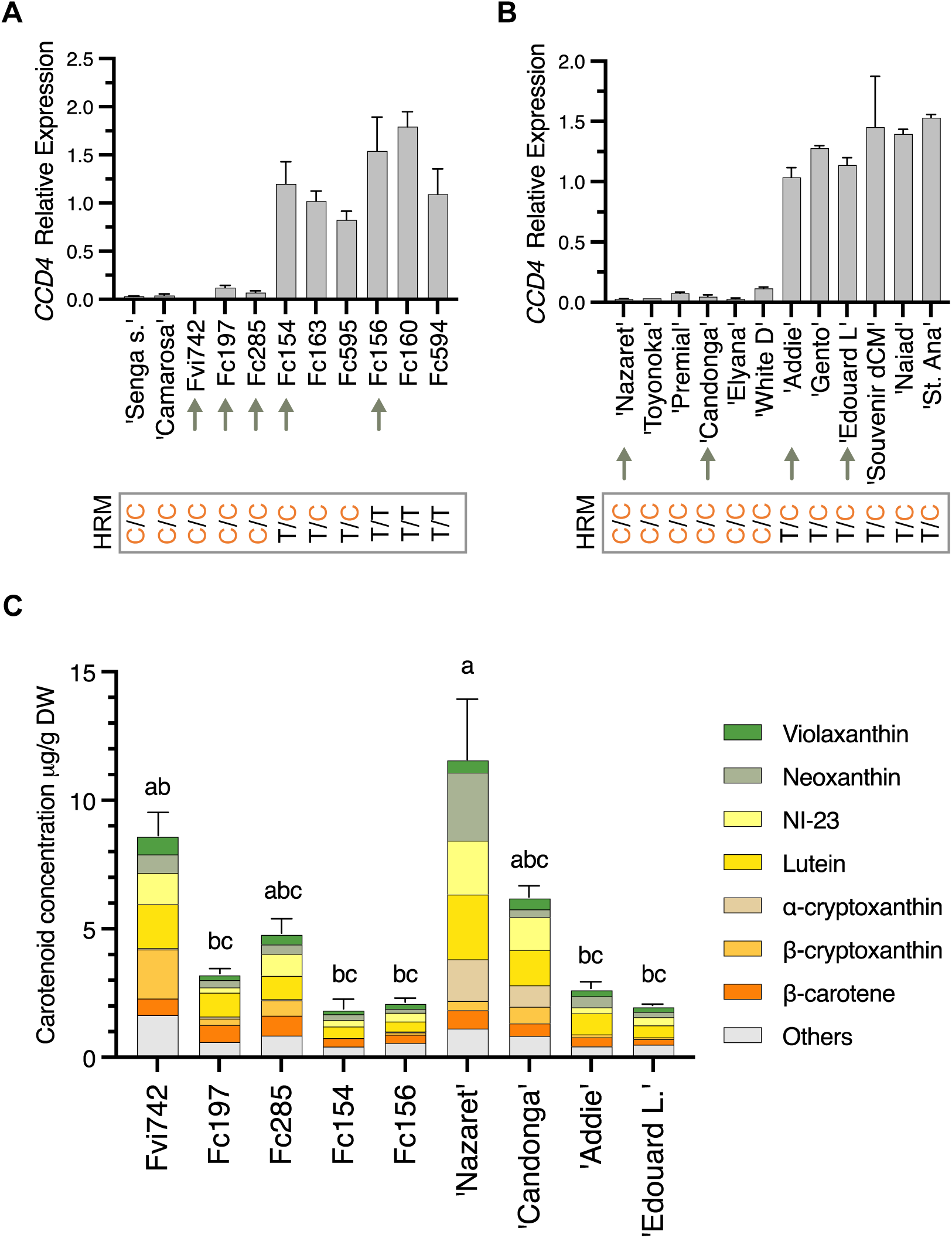
The −35 C>T SNP in *CCD4(4B)* Predicts Carotenoid Content in a Diverse Multi-species Strawberry Collection (A) −35 C>T SNP HRM genotype and *CCD4(4B)* expression in *F. virginiana* (Fvi) and *F. chiloensis* (Fc) fruits. *F.* x *ananassa* ‘Senga sengana’ and ‘Camarosa’ have been included as reference. Arrows indicate those genotypes used for carotenoid quantification in (C). (B) −35 C>T SNP HRM genotype and *CCD4(4B)* expression in a collection of *F.* x *ananassa* cultivars. Arrows indicate those genotypes used for carotenoid quantification in (C). (C) Fruit carotenoid profiles of the octoploid accessions indicated by arrows in panels (A) and (B). The total carotenoid content was averaged from three independent biological replicates, and values are presented as mean ± SEM. Each color block represents the average concentration of a specific carotenoid from the three biological samples analyzed. Statistical significance levels were determined using an ordinary one-way ANOVA followed by Tukey’s test (p < 0.05).

Carotenoid profiling in fruits from Fv742 and eight *F. chiloensis* and *F.* × *ananassa* accessions with contrasting genotypes further confirmed the negative correlation between *CCD4(4B)* expression and carotenoid accumulation (Fig. 5C). Despite notable variability in total carotenoid concentration across species and accessions, once again, lower carotenoid levels were observed in genotypes with higher *CCD4(4B)* expression, linked to the presence of the dominant *CCD4(4B)*^T^ allele. Intriguingly, in domesticated *F.* × *ananassa* accessions, selection might had favored the recessive *CCD4(4B)^C^* allele (Supplementary Table S9), suggesting a potential association of higher fruit carotenoid content with advantageous agronomic traits, a hypothesis that deserves future investigation.

Characterization of carotenoid compounds and their regulation in strawberry fruit have not received as much attention as in other fruits that accumulate larger amounts of carotenoids, such as oranges, papayas, tomato or pepper (Lado, Zacarías and Rodrigo, 2016). This work provides new insights into the genetic basis of natural variation in carotenoid content in strawberry fruits and an effective and affordable HRM marker useful as a diagnostic genetic tool to predict fruit carotenoid levels. Notably, this marker demonstrates predictive capability across plants grown in completely different environments under different cultivation management (Germany and Spain) and in multiple strawberry species, including *F. virginiana* and *F. chiloensis*, the two progenitors of cultivated *F.* × *ananassa* and sources of diversity potentially available for breeding new cultivars and the introgression of missing genes into modern varieties. Our findings will be helpful for enhancing the nutritional value of strawberry and for understanding the molecular mechanisms controlling carotenoid and apocarotenoid metabolism in this crop.

## Material and Methods

### Plant materials

The *F.* × *ananassa* ‘Senga Sengana’ × *F. chiloensis* ssp. *lucida* USA2 (SS×FcL) mapping population consists of 105 F_2_ lines. Details about population generation have been previously described (Castillejo *et al*., 2020). The population was grown under field conditions at Hansabred (Dresden, Germany) during season 2019. The parental line USA2 is a male individual that does not set fruit, and therefore, the trioecious relative USA1, which is a female and fertile plant collected from the same location, was used for phenotypic and molecular analyses.

*F. chiloensis*, *F. virginiana* and *F.* × *ananassa* accessions belong to the *Fragaria* germplasm collection at IFAPA (ESP138). They are maintained and grown in shaded greenhouses (standard or for GMOs in the case of agroinfiltrated plants) under natural sunlight and temperature conditions at the IFAPA Centre in Churriana, Málaga, Spain. All the accessions are listed in the Supplementary Table S8 (*F. chiloensis* and *F. virginiana)* and S9 (*F.* × *ananassa*).

### Phenotypic evaluation

Fruit color evaluation of the SS×FcL mapping population was performed by the naked eye following a qualitative scale. White fruits were assigned a rank score of 1, and pale-yellow fruits were scored as 2. In this study, white fruits are those lacking the yellow pigment in their flesh, but they still could accumulate anthocyanins and be red colored. Similarly, yellow fruits can also accumulate anthocyanins. Elevated anthocyanin in the flesh masks the yellow pigmentation, hindering phenotypic evaluation in 27 out of the 105 F_2_ lines (Fig. 1C).

### QTL mapping

QTL mapping was performed using the SS×FcL linkage map generated previously (Castillejo *et al*., 2020), the flesh color score as presence/absence of yellow pigmentation and the software MapQTL6 (van Ooijen, 2009 Wageningen, The Netherlands). As stated adobe, only 78 F_2_ lines were included for analysis using interval mapping (IM), restricted Multiple QTL mapping (rMQM) and the nonparametric Kruskal–Wallis test (KW) following the same pipeline as in (Castillejo *et al*., 2020). Linkage groups were named according to the proposed A-D nomenclature in (Hardigan *et al*., 2021b). The significance LOD threshold was estimated as 5 with a 1,000-permutation test. Linkage map and QTL was drawn using MapChart 2.2 (Voorrips, 2002).

Marker physical positions were inferred by BLAST against the ‘Camarosa’ reference genome (Edger et al., 2019). We searched for candidate genes within the 1-LOD QTL interval using gene annotations in the *F.* × *ananassa* ‘Camarosa’ v1. 0 genome (Edger *et al*., 2019). Linkage map with genetic distances, physical positions, genotypes and genes in the QTL interval can be found in Supplementary Tables S1-3).

### Extraction and saponification of carotenoids

Five grams of cryopreserved grinded strawberry fruit were gently mixed with 2 × 20 mL of the extracting solvent (dichloromethane/methanol/acetone, 50:25:25 v/v/v, containing 0.1% butylated hydroxytoluene) and centrifuged for 15 min at 5000 rpm. After centrifugation, the layer containing the carotenoid pigments was recovered and washed with water (2 × 5 mL) to remove any trace of acetone and methanol. Saponified carotenoids were obtained after treatment with 12 mL of methanolic KOH (30% w/v) for 1 h under dim light at room temperature. The resulting extract was washed with water (3 x 5 mL) and dried in a vacuum concentrator to evaporate the dichloromethane. The remaining solid was recovered with 500 μL of methanol/methyl tert-butyl ether (MTBE), 50:50 (v/v).

### Analysis of Carotenoids by HPLC

Chromatographic separation of carotenoids was performed with an HPLC system (Perkin Elmer series 200) equipped with a diode array detector and a C30 reverse phase column (4,6mm × 150mm 5μm). Elution conditions included two mobile phases: Methanol/water 96:4 v/v (phase A) and methyl tert-butyl ether (phase B). The elution gradient was 0 min, 95% phase A + 5% phase B; 10 min, 90% phase A + 10% phase B; 40 min, 55% phase A + 45% phase B; 45min, 25% phase A+ 75% phase B; 50 min, 100% phase B; 57min, 95% phase A + 5% phase B. The mobile phase was pumped at 0.75 mL/min, and chromatograms were monitored at 450 nm and 285 nm.

HPLC grade violaxantin, neoxanthin, lutein, zeaxanthin, phytoene, α-cryptoxanthin β-cryptoxanthin, α-carotene, β-carotene and lycopene, were used as standards for identification and quantification of carotenes and xanthophylls. Identification of carotenoids was carried out by comparison of the HPLC retention times with corresponding standards. Carotenoids were quantified using seven-pointed calibration curves that were linear (correlation coefficients ≥ 0.99) in the working range: 0.125–12.5 mg/l.

Carotenoid concentration was expressed as μg per g of dry weight (DW) of the corresponding fruit sample. DW was calculated gravimetrically by drying fruit samples at 104 °C for two days.

### Nucleic acid extraction and RT-qPCR

DNA was extracted from young leaves using the CTAB method (Doyle and Doyle, 1990)with minor modifications. Total RNA was isolated from three biological replicates following the CTAB protocol described in (Gambino, Perrone and Gribaudo, 2008) using 200 mg (leaves) or 300 mg (fruit samples) of frozen powdered samples. RT–qPCR (Reverse Transcription-Quantitative PCR) was carried out as previously described (Muñoz-Avila *et al*., 2022). Primer sequence information is listed in Supplementary Table S11.

### *CCD4(4B)* CDS and promoter cloning

The *CCD4(4B)* gene lacks introns and therefore its coding sequence (CDS) was amplified from gDNA using subgenome-specific primers ccm119 and ccm120 (Supplementary Table S1). To identify polymorphisms, the CDS was isolated from the SS×FcL parental lines (SS and USA2), four F_2_ lines with contrasting flesh color phenotypes (yellow lines F_2_-36 and 83 and white lines F_2_-01 and 52) and USA1. *CCD4(4B)* promoter sequences (*CCD4(4B)_pro_*) were amplified from the accessions listed in Supplementary Table S7 using primers ccm133 and ccm134 (Supplementary Table S1). All PCRs were performed with iProof™ High-Fidelity DNA Polymerase (Biorad) following manufacturer’s recommendations. Two ul of each PCR product were used in the ligation reaction into pCR™-Blunt cloning vector (Invitrogen™) and Sanger sequenced for polymorphisms analysis.

### Transient Agrobacterium-mediated overexpression of *CCD4(4B)* in strawberry fruits

*CCD4(4B)* CDS was PCR amplified from vector pCRBlunt-F2-52 *CCD4(4B)* with primers ccm161 and ccm88 (Supplementary Table S1) to add the Gateway™ attB1 and attB2 recombination sites. The PCR product was BP recombined into pDONR221 using Invitrogen™ Gateway™ BP Clonase™ Enzyme Mix. The resulting pDONR221-F2-52 *CCD4(4B)* was sequence validated and LR recombined (Gateway™ LR Clonase™ II) into the binary vector pK7WG2D, intended for gene overexpression driven by the cauliflower mosaic virus CaMV35S (35S) promoter (Karimi, Inzé and Depicker, 2002). *F.* × *ananassa* ‘Rociera’ fruits were coinfiltrated with *Agrobacterium tumefaciens* (*A. tumefaciens*; strain AGL0) transformed with pK7WG2D-*35S:CCD4(4B)* and *A. tumefaciens* LBA4404 harboring the p19 suppressor of gene silencing (Voinnet *et al*., 2003). A suspension of a 1:1 mix of each culture was injected in late green/early white stage fruits as previously described in (Hoffmann, Kalinowski and Schwab, 2006). As control, p19 was coinjected with a pBI-GUS vector, for expression of the GUS gene under the same 35S promoter. Fruits were harvested when reached the ripe stage for gene expression analysis and carotenoid determination. Biological replicate 1 (R-1) in control and *35S:CCD4(4B)* samples consist of a pool of three infiltrated fruits each. The remaining two replicates, R-2 and R-3, contain a single fruit each in both control and *35S:CCD4(4B)* samples.

### Agarose and HRM marker for *CCD4(4B)* genotyping

For the development of an agarose marker to detect the MITE insertion at −714 bp *CCD4(4B)_pro_*, the Geneious software (v. 7.1.9) was used to visualize all alleles from *CCD4(4B)* and the homoeologous copies from the other subgenomes. Subgenome-specific primers where designed based in the alignment produced on Geneious using Primer-BLAST (https://www.ncbi.nlm.nih.gov/tools/primer-blast/). gDNA was amplified from all assayed accessions using primers ccm155 and ccm156 (Supplementary Table S1) which flank the insertion site and amplify a 0.32 kb fragment or 0.75 kb with the MITE insertion.

For High-Resolution Melting (HRM) marker development, subgenome specific primers ccm157 and ccm158 (Supplementary Table S1) targeting the −35 C>T SNP (Position Fvb4-4:1507780 as in ‘Camarosa’ Genome Assembly v1.0.a1 (Edger *et al*., 2019)) were designed using PolyOligo 1.0 (https://github.com/MirkoLedda/polyoligo, (Ledda *et al*., 2019)). PCR amplification was performed with the Precision Melt Supermix (Bio-Rad) in a 10 μL reaction containing 0.2 μM of each primer and 20 ng of gDNA. PCR conditions were as recommended by the manufacturer, using 60°C for annealing/extension and 45 cycles of amplification. HRM variants were analyzed with the Bio-Rad Precision Melt Analysis™ Software.

### Statistical analysis

GraphPad Prism 9.4.1 was used for statistical analysis. Pairwise comparisons were performed with an unpaired t-test and multiple comparisons with one-way ANOVA followed by a post hoc Tukey’s test (P < 0.05). Statistical test employed in each case is stated in the corresponding figure legend.

## Accession Numbers

*Fragaria* × *ananassa* ‘Camarosa’ v1.0.a1:

*FaCCD4(4A)*: FxaC_13g03380; *FaCCD4(4B)*: FxaC_14g02960; *FaCCD4(4C)*: FxaC_15g02730

*Fragaria* × *ananassa* ‘Royal Royce’ v1.0:

*FaCCD4(4A)*: Fxa4Ag103306.1; *FaCCD4(4B)*: Not annotated; *FaCCD4(4C)*: Fxa4Cg202920.1

Sequence data from this article can be found in the EMBL/GenBank data libraries under accession number(s) XX000000

## Supplementary data

The following materials are available in the online version of this article.

Supplementary Fig. S1. Expression of *CCD4* homoeologs in ‘Camarosa’ tissues.

Supplementary Fig. S2. *CCD4(4B)* promoter alignment of SSxFcL population and other contrasting *F. chiloensis* and *F*. x *ananassa* accessions.

Supplementary Fig. S3. Genotyping of the SS×FcL mapping population using the marker to detect the MITE insertion in the CCD4(4B) promoter region.

Supplementary Table S1. Linkage map and genotypes in the SS×FcL population.

Supplementary Table S2. Candidate genes in the qYellow Flesh-4B QTL interval in ‘Camarosa’.

Supplementary Table S3. Candidate genes in the qYellow Flesh-4B QTL interval in *F. vesca*.

Supplementary Table S4. Phenotype and HRM (−35 C>T SNP) genotype in the SS×FcL mapping population.

Supplementary Table S5. Fruit carotenoid content in the SS×FcL population.

Supplementary Table S6. Fruit carotenoid content in each biological replicate from the *35S:CCD4(4B)* transient expression assay.

Supplementary Table S7. *CCD4(4B)* promoter polymorphisms associated with differential *CCD4(4B)* expression level in *Fragaria* accessions.

Supplementary Table S8. HRM (−35 C>T SNP) genotype in *F. chiloensis* and *F. virginiana* accessions.

Supplementary Table S9. HRM genotype (−35 C>T SNP) in the *F.* × *ananassa* cultivars and hybrids collection

Supplementary Table S10. Fruit carotenoid content in the octoploid *Fragaria* spp. Collection

Supplementary Table S11. Primers used in this work.

## Author Contributions

C.C. and I.A. designed and performed the research, analyzed the data and wrote the paper with input from all other authors; F.J. R.-G., J.L. O.-D. and J.M. M.-R. conducted carotenoid quantification; H.W., V.W. and K.O. developed the octoploid SS×FcL mapping population and contributed to the phenotypic analysis. R.T. performed the genotyping of the natural population with the HRM marker. J.F.S.-S. contributed diverse plant material, phenotyping and analyzed sequence data.

## Funding

This work was supported by Ministerio de Ciencia e Innovación and Agencia Estatal de Investigación (PID2019-111496RR-I00 and PID2022-138290OR-I00 / MCIN/AEI / 10.13039/501100011033 / FEDER), the European Union’s H2020 research and innovation programme (BreedingValue project, grant agreement No 101000747) and by the German Federal Ministry of Education and Research (BMBF, FKZ 031A216 A and B). The *Fragaria* collection at IFAPA is financed by IFAPA Project PR.CRF.CRF202200.002 with funds from the European Agricultural Fund for Rural Development.

## Acknowledgments

We’re thankful to Francisco J. Durán at IFAPA for the meticulous care and maintenance of the strawberry plants.

## Conflict of interest statement

The authors declare no conflict of interest

---

**Supplementary Fig. S1.**
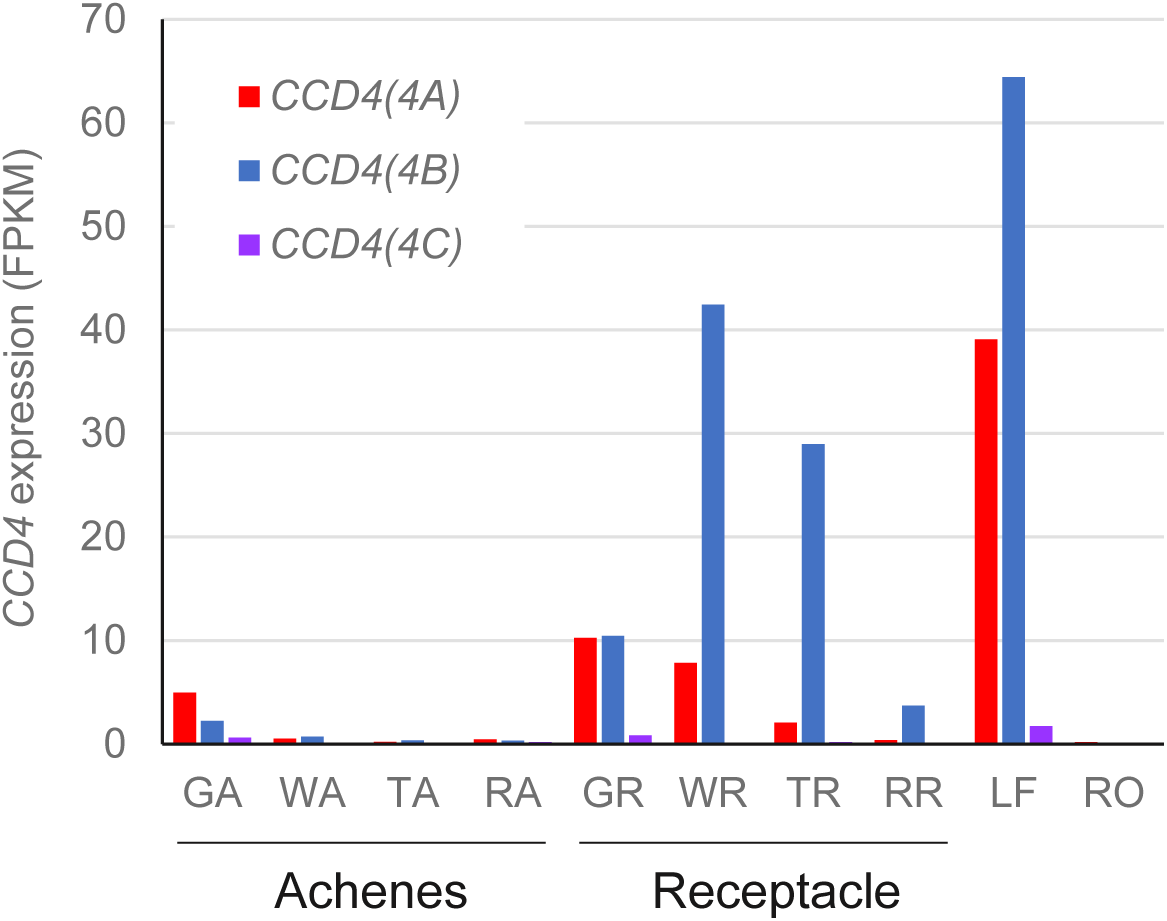
(Supports Fig. 1) Expression of *CCD4* homoeologs from a RNA-seq experiment of ‘Camarosa’ tissues. Expression level dominance is biased towards *CCD4(4B)*. GA: Green Achene; WA: White Achene; TA: Turning Achene; RA: Red Achene; GR: Green Receptacle; WR: White Receptacle; TR: Turning Receptacle; RR: Red Receptacle; LF: Leaf; RO: Root. Gene codes in the ‘Camarosa’ genome: *CCD4(4A): FxaC_13g03380; CCD4(4B): FxaC_14g02960; CCD4(4C): FxaC_15g02730*

**Supplementary Figure S2.**
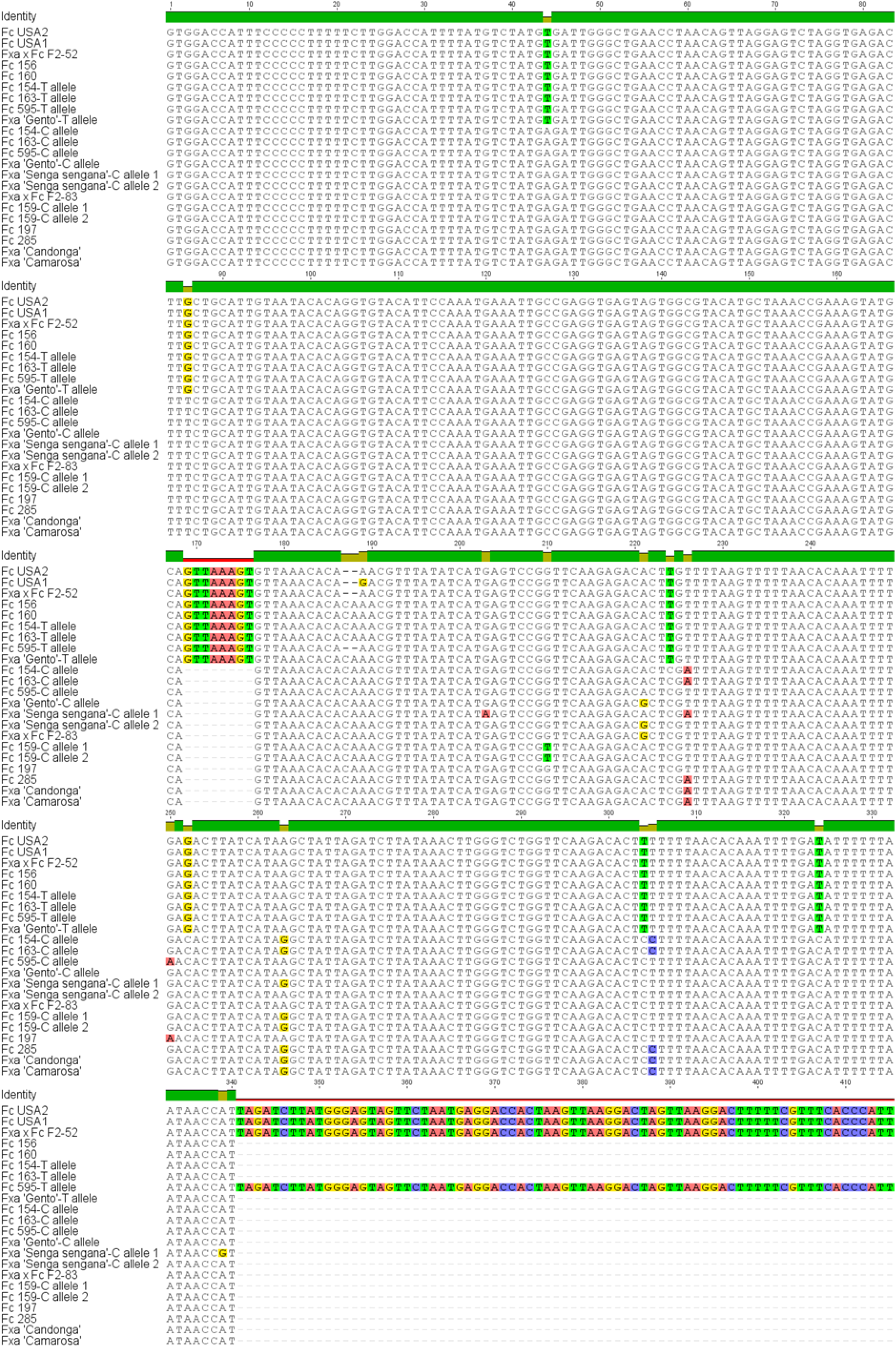
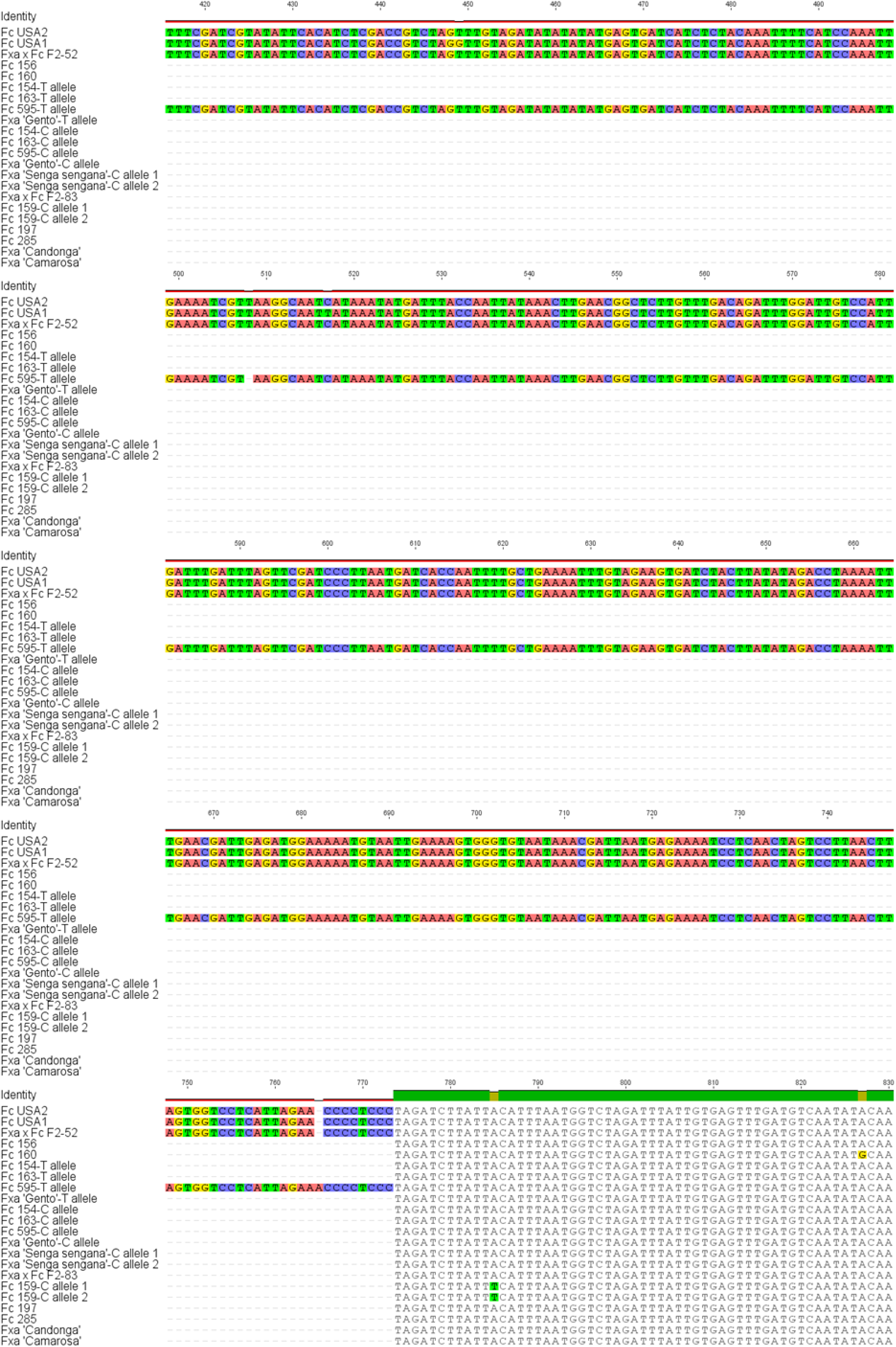
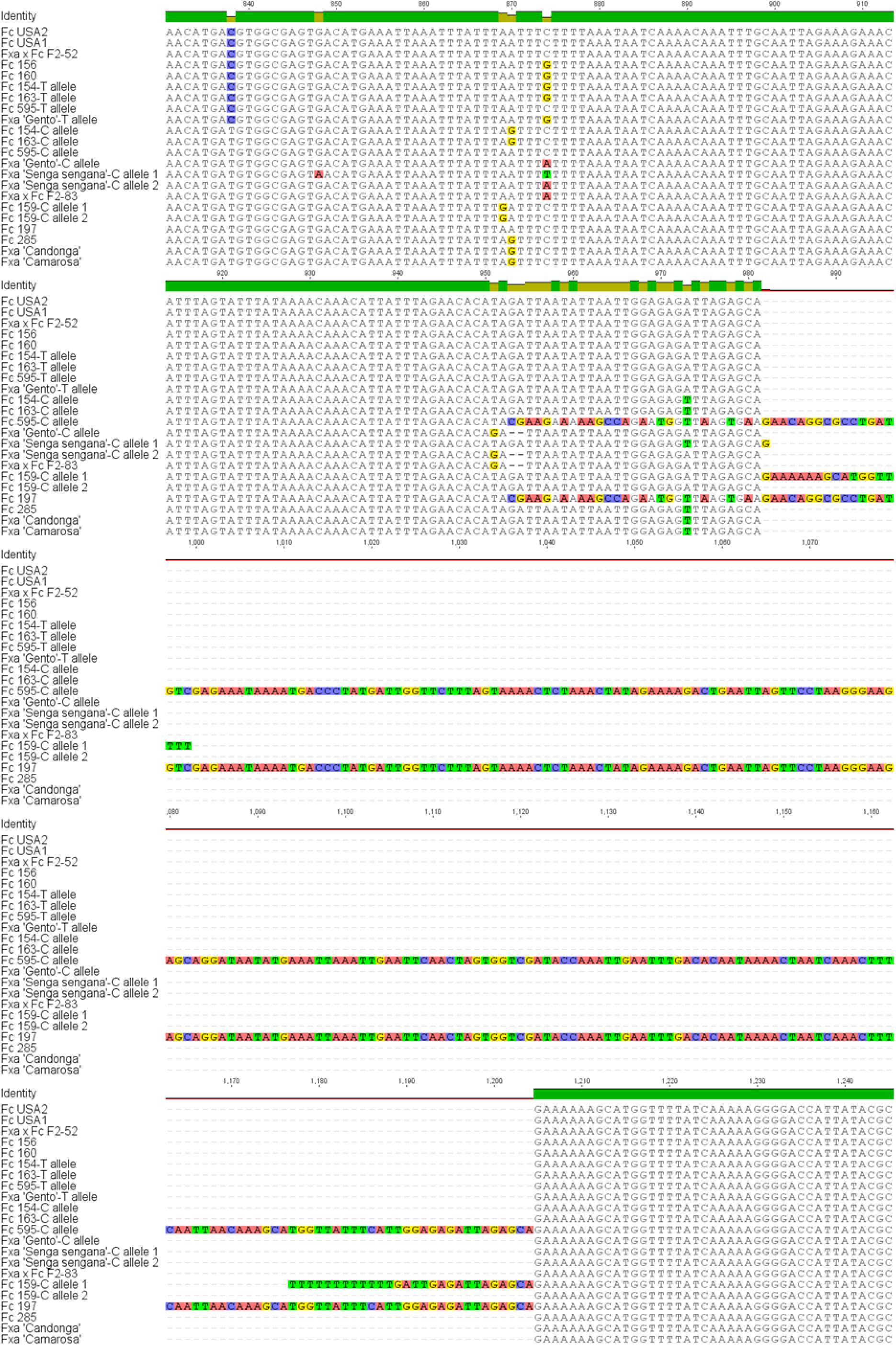
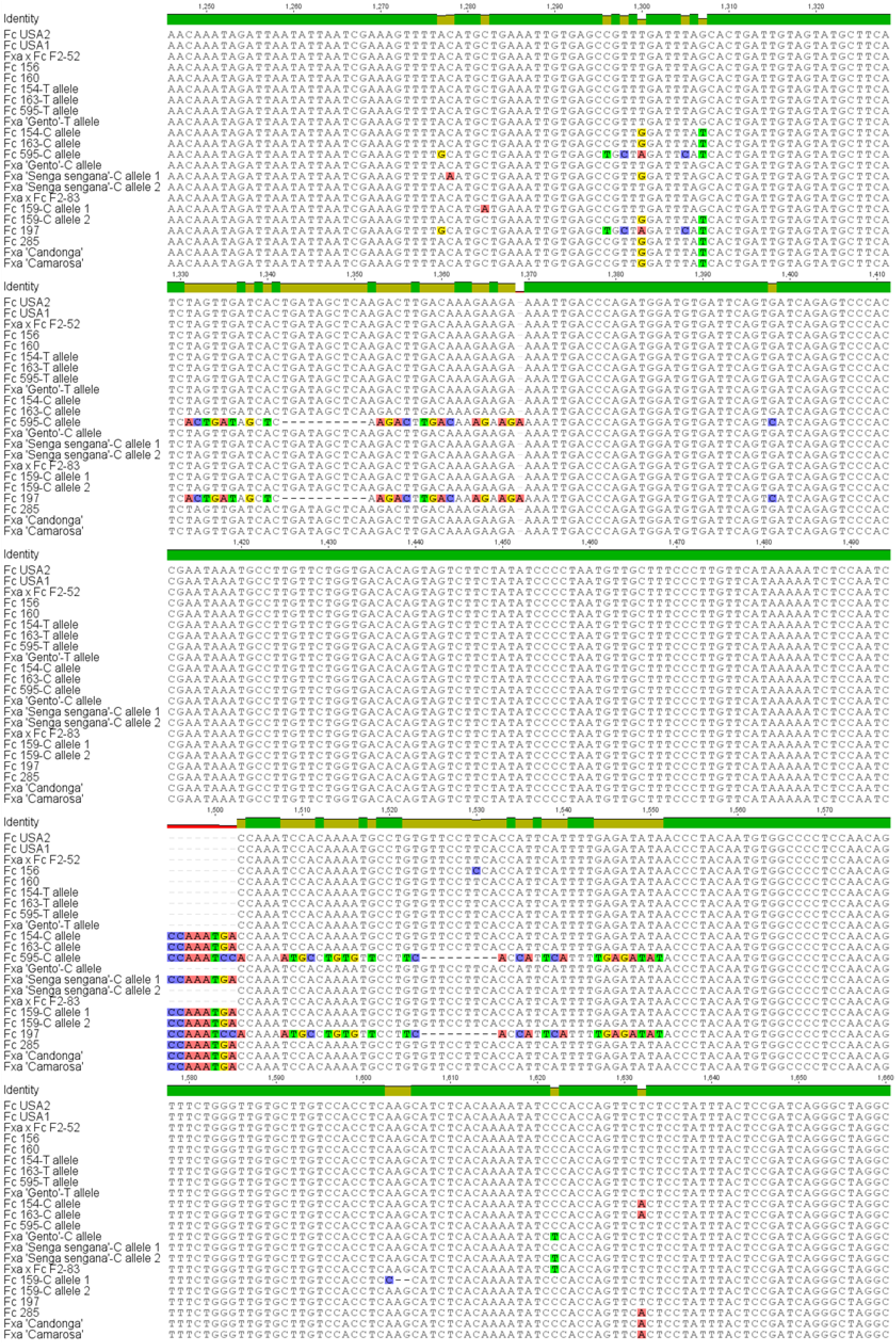
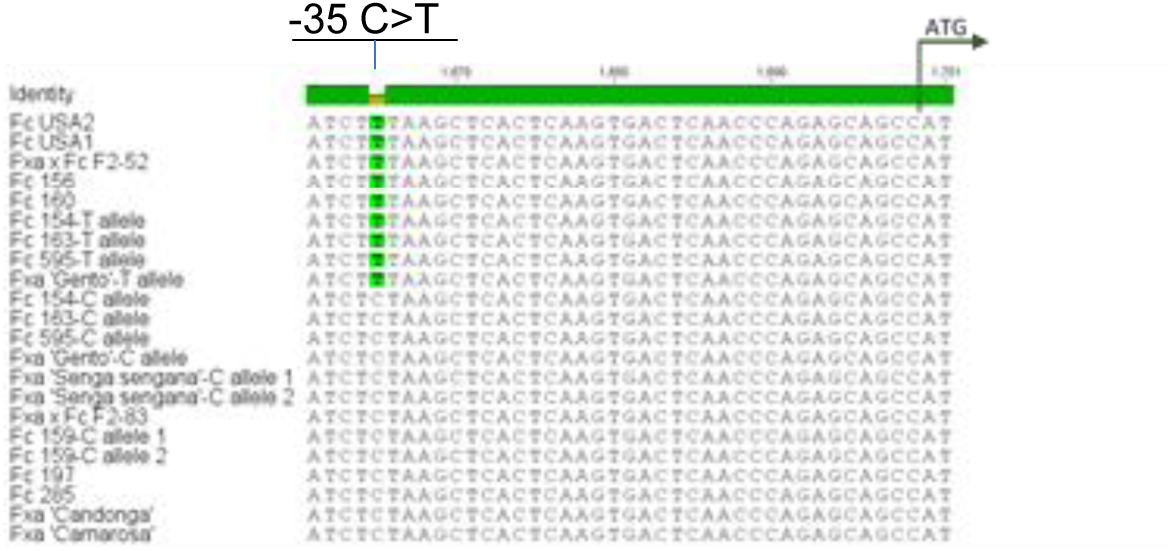
(Supports Fig. 2) CCD4(4A) promoter alignment of SSxFcL population and other contrasting *F. chiloensis* and *F.* x *ananassa* accessions. *CCD4(4B)* promoter alignment from the two parental of the SSxFcL population: *Fragaria* x *ananassa* (Fxa) ‘Senga sengana’ and *Fragaria chiloensis* (Fc) subsp. lucida USA2 (male). USA1 is the female relative of USA2. Two contrasting F_2_ siblings are also shown (white fleshed F_2_-52 and yellow fleshed F_2_-83). When non homozygous, the two alleles were included in the alignment. T/C allele makes reference to the nucleotide at −35 C>T SNP. White fleshed accessions: Fxa ‘Gento’, Fc USA2, Fc USA1, Fc 156, Fc 160, Fc154, Fc 163 and Fc 595. Yellow fleshed accessions: Fxa ‘Camarosa’, Fxa ‘Candonga’, Fxa ‘Senga sengana’, Fc 159, Fc197, Fc 285.

**Supplementary Fig. S3.**
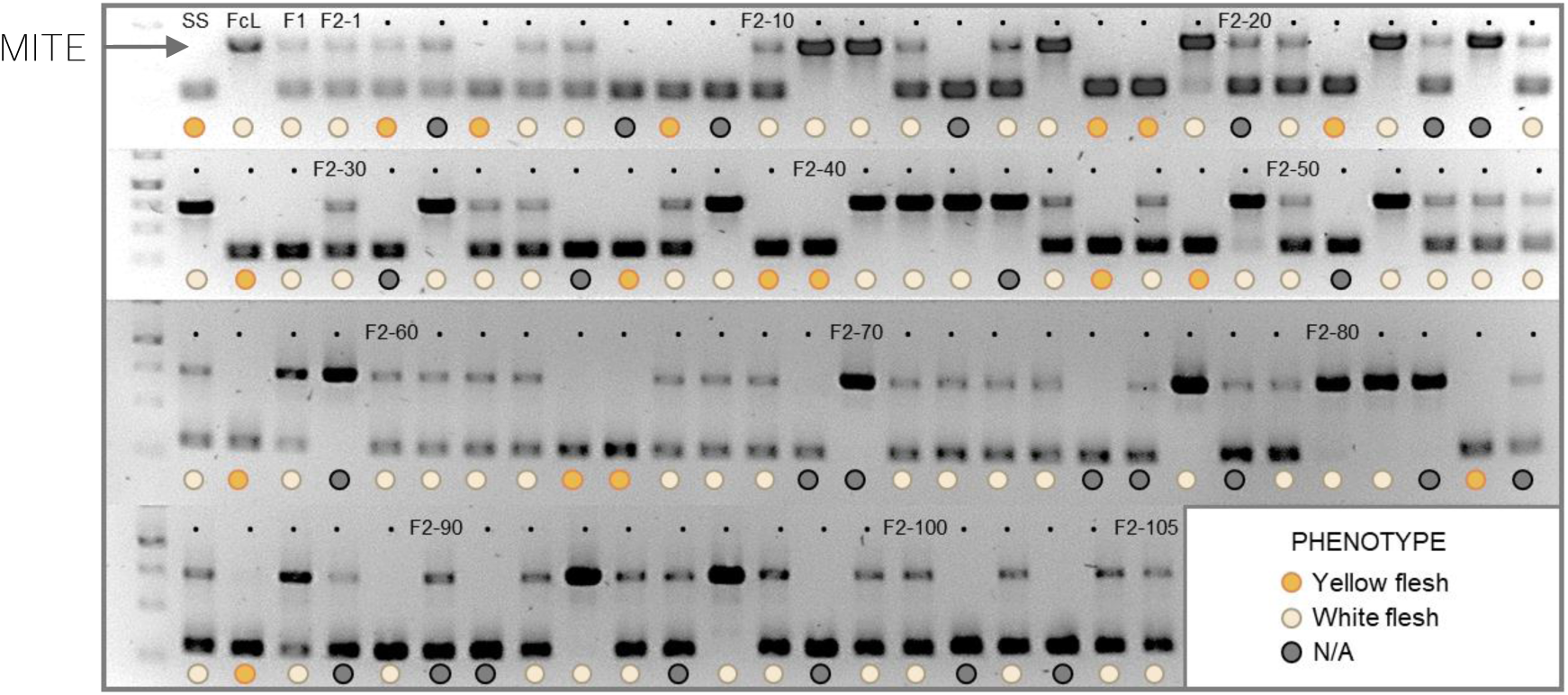
(Supports Fig. 3) Genotyping of the SS×FcL mapping population using the marker to detect the MITE insertion in *CCD4(4B)* promoter region. Primers used in the PCR flank the MITE insertion site. Upper band corresponds to the promoter fragment harboring the MITE insertion. Flesh color phenotype of each individual is represented with a color code (see legend). Individuals F_2_-3, w/o fruits, and those with high anthocyanin accumulation could not be rated for white/yellow hue (N/A). In all cases, except for individuals F_2_-2 and F_2_-29, the presence of the transposon insertion was associated with white flesh.

## Notes

### Competing Interest Statement

The authors have declared no competing interest.

